# An oxadiazole-based compound potentiates anti-tuberculosis treatment by increasing host resistance via zinc poisoning

**DOI:** 10.1101/2023.07.21.549986

**Authors:** Alexandra Maure, Emeline Lawarée, Francesco Fiorentino, Alexandre Pawlik, Saideep Gona, Alexandre Giraud-Gatineau, Matthew J.G. Eldridge, Anne Danckaert, David Hardy, Wafa Frigui, Camille Keck, Nathalie Aulner, Antonello Mai, Mélanie Hamon, Luis Barreiro, Priscille Brodin, Roland Brosch, Dante Rotili, Ludovic Tailleux

**Affiliations:** Institut Pasteur, Université Paris Cité, CNRS UMR 6047, Unit for Integrated Mycobacterial Pathogenomics, F-75015 Paris, France; Department of Drug Chemistry and Technologies, Sapienza University of Rome, 00185 Rome, Italy; Department of Genetic Medicine, University of Chicago, Chicago, USA; Institut Pasteur, Université Paris Cité, CNRS UMR 6047, Biology of Spirochetes Unit, F-75015 Paris, France; Institut Pasteur, Université Paris Cité, Chromatine et Infection unit, F-75015 Paris, France; Institut Pasteur, Université Paris Cité, UTechS BioImaging-C2RT, F-75015 Paris, France; Institut Pasteur, Université Paris Cité, Histopathology Platform, F-75015 Paris, France; Pasteur Institute, Cenci-bolognetti Foundation, Sapienza University of Rome, 00185 Rome, Italy; Université de Lille, CNRS, INSERM, CHU Lille, Institut Pasteur de Lille, U1019 - UMR 8204 - CIIL - Center for Infection and Immunity of Lille, F-59000 Lille, France

**Keywords:** tuberculosis, macrophage, host-directed therapy, zinc

## Abstract

Anti-tuberculosis drugs, mostly developed over 60 years ago, combined with a poorly effective vaccine, have failed to eradicate tuberculosis. More worryingly, multi-resistant strains of *Mycobacterium tuberculosis* are constantly emerging. Innovative strategies are thus urgently needed to improve tuberculosis treatment. Recently, host-directed therapy has emerged as a promising strategy to be used in adjunct with existing or future antibiotics, by improving innate immunity or limiting immunopathology. Here, using high content imaging, we identified novel 1,2,4-oxadiazole-based compounds, that allow human macrophages to control MTB replication. Genome-wide gene expression analysis revealed that these molecules induced zinc remobilization inside cells, resulting in bacterial zinc intoxication. More importantly, we also demonstrated that, upon treatment with these novel compounds, *M. tuberculosis* became even more sensitive to anti-tuberculosis drugs, *in vitro* and *in vivo*, in a mouse model of tuberculosis. Manipulation of heavy metal homeostasis holds thus great promise to be exploited to develop host-directed therapeutic interventions.

## Introduction

Antibiotics have considerably extended human’s life span by changing the outcome of bacterial infections. With this medical breakthrough came the emergence of antimicrobial resistance worldwide, representing one of the biggest threats to human health. Tuberculosis (TB) is no exception, with the clock ticking for TB care and prevention. TB is a disease caused by the bacterium *Mycobacterium tuberculosis* (MTB), that is spread from person to person through the air. Although significant progress has been made to tackle TB over the last few decades, with the discovery of major anti-tuberculosis drugs (1), the disease still kills over 1.6 million people annually. Worryingly, multidrug resistant (MDR) strains of MTB, which are resistant to both the front-line anti-TB drugs rifampicin (RIF) and isoniazid (INH), have emerged and are widespread worldwide (2). In 2021, 450,000 patients were diagnosed with rifampicin-resistant TB (RR-TB) or MDR-TB (2); 20% of those patients also show resistance to fluoroquinolones, an important drug class for the treatment of drug-resistant TB (2). Typically, drug-sensitive TB can be cured by a 6-month treatment, combining up to 4 antibiotics, namely INH, RIF, ethambutol (EMB) and pyrazinamide (PZA). Curing MDR-TB is more difficult and requires treatments of up to 2 years with more toxic drugs, often with limited success. While new antibiotics are being developed and brought to the clinic, resistance to these molecules are rapidly detected. New strategies are thus urgently needed to prevent the emergence of drug resistance and to shorten treatment duration.

Host-directed approaches are a promising strategy to be used in adjunct with existing or future antibiotics (3, 4). Several host-directed therapy (HDT) to treat TB have been developed and mostly act in (i) the manipulation of host anti-mycobacterial pathways that are blocked or altered by MTB to promote its survival, (ii) the potentiation of antimicrobial host immune defense mechanisms or (iii) the amelioration of immunopathology (4). Macrophages represent a prime target for HDT as they play a key role in the outcome of a mycobacterial infection. On one hand, they orchestrate the formation of granulomas, present mycobacterial antigens to T cells and can kill the bacillus upon IFN-γ activation (5). One the other hand, macrophages are the primary host niche for MTB, which has developed different strategies to survive and to multiply inside the macrophages’ phagosome. These include prevention of phagosome acidification (6), inhibition of phagolysosomal fusion (7) and phagosomal rupture (8, 9).

Some *in vitro* studies have demonstrated that HDT-compounds are indeed effective at increasing macrophage resistance to MTB infection (3). These molecules include repurposed FDA-approved drugs or new compounds that have been identified by high-throughput screening of MTB-infected cells. Most of them promote phagosome maturation (e.g., tyrosine kinase inhibitors), activate autophagy (such as inhibitors of the mechanistic target of rapamycin (mTOR)), induce antimicrobial peptides or inhibit lipid body formation (3). Here, we identified new molecules able to limit MTB growth in human macrophages by inducing zinc remobilization inside cells, resulting in bacterial poisoning. Interestingly, these compounds also potentiate the activity of known anti-TB drugs not only *in vitro* but also *in vivo* in a TB mouse model.

## Results

### Identification of a compound that inhibits the intracellular MTB growth

We and others have previously shown that numerous epigenetic modifications occur in myeloid cells upon MTB infection (10–12). These changes are either part of the initiation of the host response or an immune escape mechanism from MTB. MTB indeed manipulates epigenetic host-signaling pathways to subvert host immunity (10). Reprogramming the host immune system by a compound targeting the host epigenome may thus lead to a better control of the bacterial infection (13). To tackle this hypothesis, we screened an in-house library of 157 epigenetic compounds in MTB-infected macrophages using high-content imaging. This library comprises activators and inhibitors of enzymes which carry out various epigenetic modifications. For instance, it includes molecules that modulate the activity of DNA methyltransferases (DNMTs), histone methyltransferases (HMTs) and demethylases (HDMs), histone acetyltransferases (HATs) and deacetylases (HDACs) and poly (ADP-ribose) polymerases (PARPs) (S1 Table).

Briefly, human monocyte-derived macrophages were infected with MTB expressing GFP at a MOI of 0.5, before being seeded in a 384-well plate; each well contained a different compound at a final concentration of 10 μM. After 96 h of infection, cells were fixed and nuclei were stained with Hoechst 33342. Fluorescence was analyzed by automated confocal microscopy. Cell toxicity was determined by comparing the number of nuclei stained by Hoechst 33342 in treated wells with the negative control well (DMSO). Only compounds associated with a cell viability greater than 75% were considered for further analysis. 106 molecules were non-toxic to human macrophages (Fig 1A). Inhibition of intracellular MTB growth in presence of the remaining compounds was then assessed by determining the GFP area per cell, which is correlated to the amount of intracellular MTB. We identified one molecule, MC3465, which reduces the bacterial growth by 60% compared to the control (Fig 1A and 1B). This result was confirmed by enumerating the number of bacteria inside macrophages treated with MC3465 for 24 h and 96 h. A 25% and 55% decrease of colony-forming units (CFUs) was observed after 24 h and 96 h of treatment compared to the DMSO control, respectively (Fig 1C). There are substantial differences in the virulence of MTB strains. These differences could lead to an attenuated activity of our compound. We thus tested the efficacy of MC3465 in macrophages infected with highly-virulent clinical isolates of MTB, namely CDC1551, GC1237, and Myc5750 (the last two belonging to the Beijing family) (14–16). After 96 h treatment, we observed a substantial decrease in bacterial survival rates with MC3465 (Fig 1D). This compound did not show any cell toxicity, even at higher concentration (50 µM) over an incubation period of 7 days (S1A Fig). MC3465 was equally active in macrophages differentiated in the presence of GM-CSF or M-CSF, two cytokines known to prime toward pro-inflammatory (M1) and alternatively activated (M2) phenotypes (17) (S1B Fig).

**Fig 1.**
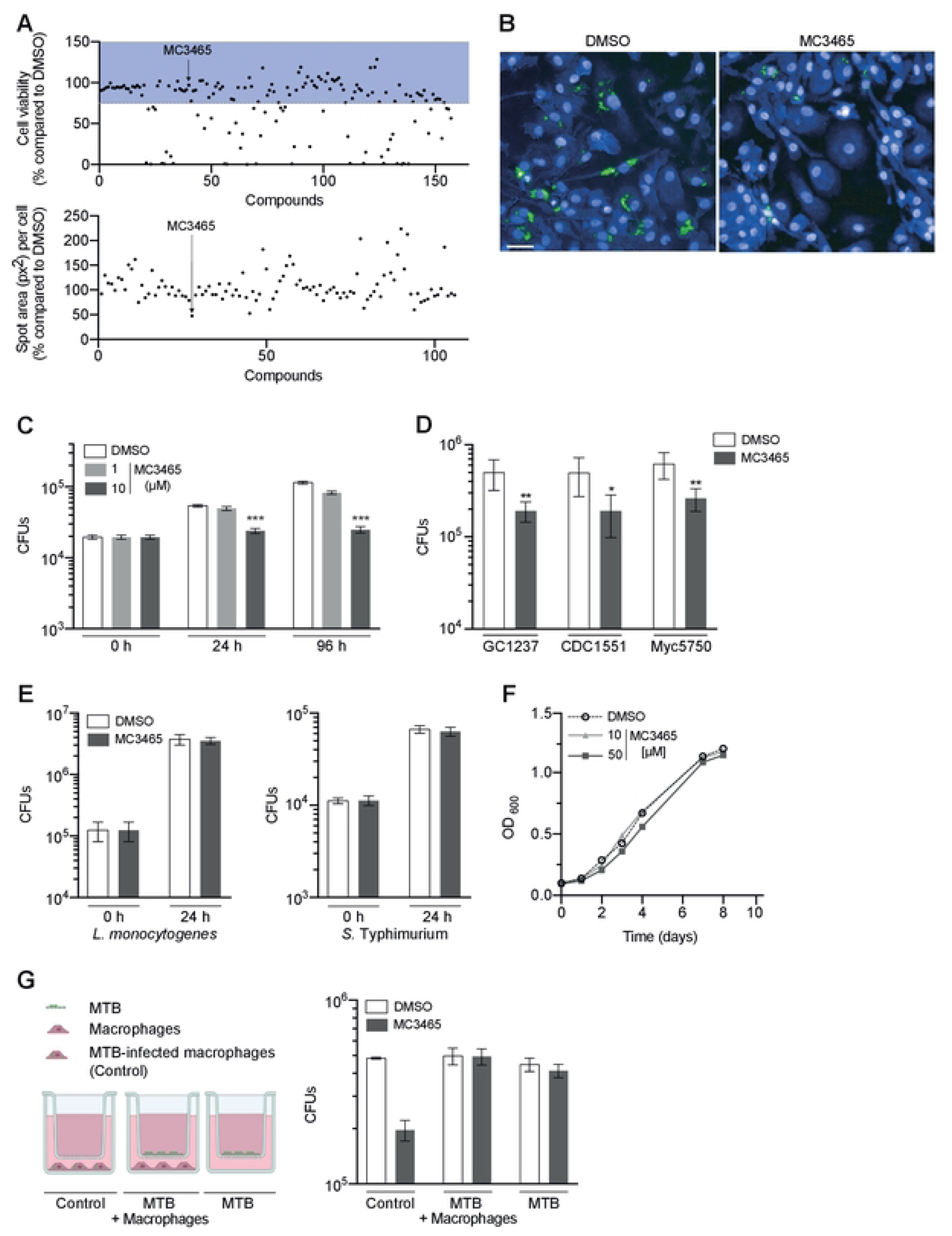
Identification of new compounds inhibiting the intracellular growth of MTB. **(A)** Human monocyte-derived macrophages were infected with GFP expressing MTB (GFP-MTB) and incubated with epigenetics-related compounds. After 96 h treatment, cells were labelled with Hoechst 33342 and HCS CellMask Blue. Fluorescence was analyzed by the Opera Phenix® Plus High-Content Screening System. Quantification of GFP staining and enumeration of cells were performed using Columbus image analysis software. Only compounds which were not toxic (cell viability>75%, in blue in the upper panel) were kept for further analysis. The number of live cells and the areas of intracellular bacteria (px: pixels, lower panel) were expressed as the percentage in compound-treated cells compared to cells incubated with DMSO. **(B)** Representative confocal images of macrophages infected with MTB (green) and treated with MC3465. Hoechst 33342 and HCS CellMask (blue) were used to delimit nuclei and cytoplasm shapes, respectively. Scale bar: 10 μm. **(C)** MTB-infected macrophages were treated with MC3465 (1 and 10 µM). The number of intracellular bacteria was enumerated at 0, 24 and 96 h post-infection. **(D)** Macrophages were infected with clinical strains of MTB, namely GC1237, CDC1551 and Myc5750 and were treated with MC3465 (10 µM). The number of intracellular bacteria was enumerated at 96 h post-treatment. **(E)** Macrophages were infected with *Listeria monocytogenes* or *Salmonella* Typhimurium. Intracellular bacteria were enumerated at 0 and 24 h post-infection. **(F)** Growth of MTB in culture liquid medium in the presence of MC3465 (10 and 50 µM). **(G)** MTB-infected macrophages, MTB separated from macrophages using Transwell® inserts and MTB alone were treated with MC3465 (10 µM) or DMSO for 96 h. Cells were lysed and the number of intracellular bacteria enumerated. One representative experiment (of at least three) is shown. Error bars represent the mean ± SD. One-way ANOVA test and t-test were used. * p < 0.05, ** p < 0.01, *** p < 0.001, ****p<0.0001.

We next tested whether MC3465 also inhibited the intracellular bacterial growth of other bacteria. Macrophages were infected with *Listeria monocytogenes* and *Salmonella enterica serovar* Typhimurium for 24 h, in presence of MC3465 (10 μM) or with DMSO alone. Cells were then lysed and CFUs were enumerated. Our results show that MC3465 does not impede the intracellular growth of *L. monocytogenes* and *S.* Typhimurium, indicating that the action of this compound could be specific to MTB-infected cells (Fig 1E).

MC3465 is supposed to act as an inhibitor of the human NAD^+^-dependent deacetylase sirtuin 2 (SIRT2) (18). However, it could have a direct inhibitory effect on the bacteria. To verify the host-directed effect of this molecule, we monitored the MTB growth in liquid medium by measuring the optical density (OD600nm). Even at a higher concentration (50 μM), MC3465 does not affect MTB growth (Fig 1F). To further confirm that MC3465 targets only intracellular MTB, bacteria were cultivated (i) within macrophages, (ii) in presence of macrophages, without any direct contact with the host cells or (iii) alone. After 96 h treatment, the number of bacteria was enumerated in each condition. As shown in Fig 1G, MC3465 only limits the multiplication of intracellular MTB. This observation suggests that MC3465 is not metabolized by macrophages in an active form that could target extracellular bacteria.

### MC3465 inhibits MTB growth in a SIRT2-independent manner

MC3465 or 3-(4-bromophenyl)-5-(3-bromopropyl)-1,2,4-oxadiazole is described to be a specific human SIRT2 inhibitor, with no activity against SIRT1, −3, and −5 (18). Recently, it has been published that the expression of SIRT2 is upregulated upon MTB infection and that inhibition of SIRT2 by AGK2 leads to a decrease of intracellular MTB survival in a human monocytic/macrophage cell line (THP-1) and in murine peritoneal macrophages (19). To assess if MC3465 limits MTB intracellular growth through SIRT2 inhibition, we incubated MTB-infected macrophages with either DMSO, MC3465 or with the two commonly used human SIRT2 inhibitors: AGK2 and SirReal2 (20). 96 h post-infection, cells were lysed and the number of bacteria was determined by CFUs. While MC3465 decreased the number of viable intracellular MTB in a dose-dependent manner, AGK2 and SirReal2 had no effect on bacterial growth, suggesting MC3465 acts in a different way to the SIRT2 inhibitors (S2A Fig). To keep investigating if SIRT2 is the putative target of MC3465, we inactivated SIRT2 expression in human macrophages using siRNA-mediated gene silencing (21). The level of SIRT2 expression decreased by about 70% upon silencing (S2B Fig). In SIRT2-silenced cells, MC3465 restrained the intracellular multiplication of MTB as well as in cells expressing normal level of SIRT2 (S2C Fig). Moreover, MC3465 was still active on intracellular MTB in mouse SIRT2^-/-^ embryonic fibroblasts (MEFs), to a similar level as the wild-type cells (S2D Fig). Altogether, our results strongly demonstrate that SIRT2 is not the target of MC3465. They also argue against a role of SIRT2 in the interactions between MTB and human macrophages.

Sirtuins are a family of NAD^+^-dependent lysine deacylases highly conserved from bacteria to humans. MTB has a NAD^+^-dependent protein deacetylase, namely Rv1151c (22). While Rv1151c is a non-essential gene for *in vitro* growth of MTB (23), we cannot exclude a role in macrophage parasitism. We generated a deletion of mutant of Rv*1151c* in H37Rv (ΔRv*1151c*). As expected, there was no difference in growth between ΔRv*1151c* and the WT MTB strain (S2E Fig). We then infected macrophages with both strains and treated the cells with MC3465. After 96 h of infection, the cells were lysed and the number of intracellular bacteria was enumerated. ΔRv*1151c* was able to grow intracellularly at a similar level as the WT strain and was as susceptible to MC3465 as the WT (S2F Fig). This result excludes an inhibition of the mycobacterial SIRT2-like protein by MC3465.

### MC3465 and its analog MC3466 alters zinc homeostasis, resulting in bacterial poisoning

Autophagy is a well-established key factor for host defense against MTB (24) and several autophagy activating drugs have been used to restrict MTB survival (3). We therefore tested whether MC3465 activated the formation of autophagosomes and autolysosomes. MTB-infected macrophages were stained with an antibody against microtubule-associated protein light-chain 3B (LC3B) and the fluorescence was analyzed by confocal microscopy. We observed minimal difference in the number of LC3B puncta per cell at 24 h and 48 h post-treatment (S3A and S3B Fig). To confirm this data, we used well-characterized autophagy inhibitors such as 3-methyladenine (3-MA), bafilomycin (BAF) or chloroquine (CQ) and tested their effect on MTB-infected macrophages treated with MC3465. 96 h post-infection, intracellular bacteria were collected and plated to enumerate the CFUs. Our data show that autophagy inhibitors 3-MA, BAF or CQ do not inhibit the effect of MC3465 (S2C Fig). MC3465 activity is thus not related to an induction of autophagy.

To further understand the molecular mechanism of action of MC3465 to restrain intracellular MTB growth, we performed a transcriptome analysis upon MC3465 treatment. We infected macrophages from four healthy donors with MTB and then treated them with MC3465. After 4 h and 24 h treatment, we characterized the genome-wide gene expression profiles of macrophages by RNAseq, with DMSO-treated cells serving as a control. The expression of 181 genes was differentially regulated following MC3465 treatment (p-value < 0.05, log FC ≥ 0.5 and ≤ −0.5) after 4 h, with 53 being upregulated and 128 being downregulated (Fig 2A). The expression of more genes was affected after 24h treatment. 224 genes were upregulated and 652 genes downregulated. (p value < 0.05, log FC ≥ 0.5 and ≤ −0.5) (Fig 2A).

**Fig 2.**
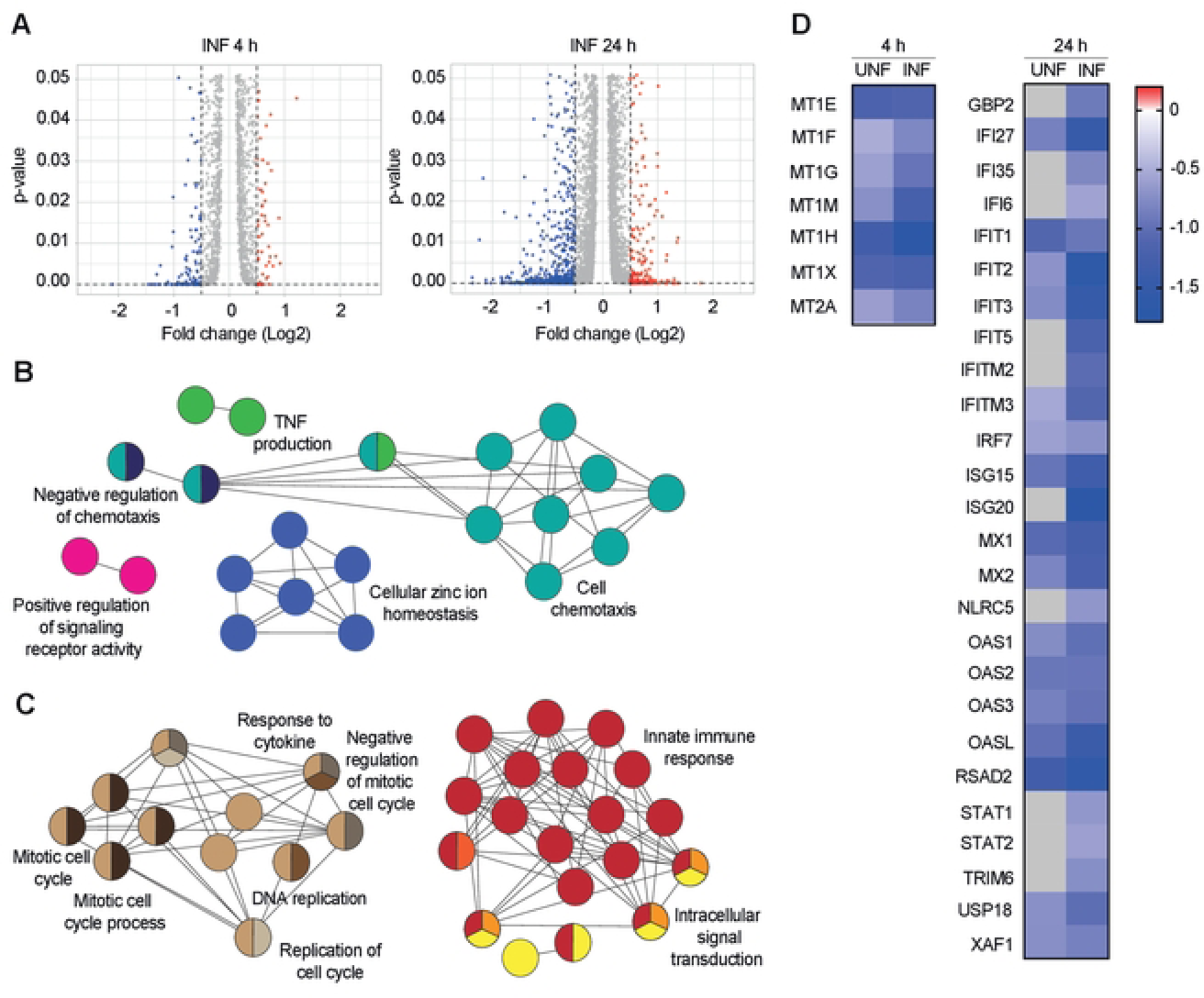
Differentially expressed genes upon MC3465 treatment. Naïve and MTB-infected macrophages derived from four individual donors were treated with MC3465 for 4 h and 24 h. Differentially expressed genes were identified by mRNAseq. **(A)** Volcano plot showing differentially expressed genes due to MC3465 treatment (p-value <0.05, Log Fold Change (FC) <-0.5 and >0.5). **(B-C)** Gene ontology enrichment analysis of genes whose expression is downregulated by MC3465 treatment in MTB-infected cells, using the Cytoscape app ClueGO (p.value<0,05; LogFC<-0,5 or LogFC>0,5) after 4 h **(B)** and 24 h **(C**). **(D)** Heatmap showing expression of genes differentially expressed in naive and MTB-infected cells upon MC3465 treatment. Genes related to cellular zinc ion homeostasis and type I interferon signaling pathway were represented at 4 and 24 h, respectively. Genes that were not differentially expressed were represented by a grey square.

We classified the differentially expressed genes by performing gene-set enrichment analysis using ClueGO cluster analysis (25). The gene set downregulated by MC3465 at 4 h was significantly enriched for genes associated with chemotaxis and cellular zinc ion homeostasis and cell chemotaxis (Fig 2B-D). At 24 h, most of the downregulated genes belongs to the innate immune response and the cytokine response, with many genes belonging to the type I IFN pathway (Fig 2C and 2D).

Maintaining intracellular zinc homeostasis is essential for all domains of life, as zinc serves as cofactor for enzymes and other proteins involved in different fundamental cellular processes. In this context, nutritional immunity is a well-known mechanism deployed by the host cell to limit the availability of essential metals to starve pathogens and hence impedes the intracellular growth of the pathogens (26). In contrast, intracellular bacteria can be intoxicated by an excess of free metal ions, namely zinc and copper (27, 28). Our transcriptomic data in MTB-infected and naïve macrophages suggested that MC3465 alters zinc sequestration. The expression of genes coding for metallothioneins was down-regulated upon treatment (Fig. 2D). Metallothioneins are cysteine-rich metal-binding proteins that are important for zinc and copper homeostasis, protection against oxidative stress, DNA damage and heavy metals toxicity (29). Their down-regulation in treated cells could therefore be associated with an increase of intracellular free zinc.

To test this hypothesis, naïve and MTB-infected macrophages were treated with MC3465 during 4 h and stained with Fluozin^TM^ 3-AM, a fluorescent Zn^2+^-selective indicator (30, 31). Cells treated with the zinc-chelating cell-permeant agent TPEN (27) were used as control. Confocal microscopy showed that the treatment resulted in a rapid increase in the Fluozin^TM^ 3-AM signal, whether or not the cells were infected (Fig 3A and 3B and S4A and S4B Fig). The labeling was as strong as in cells cultured in medium supplemented with high concentrations of ZnSO_4_ (S4B Fig). Moreover, upon treatment with MC3465, we also found numerous colocalization between the Fluozin^TM^ 3-AM signal and DsRed-expressing MTB, indicating a zinc accumulation in bacteria-containing phagosomes (Fig 3C and 3D).

**Fig 3.**
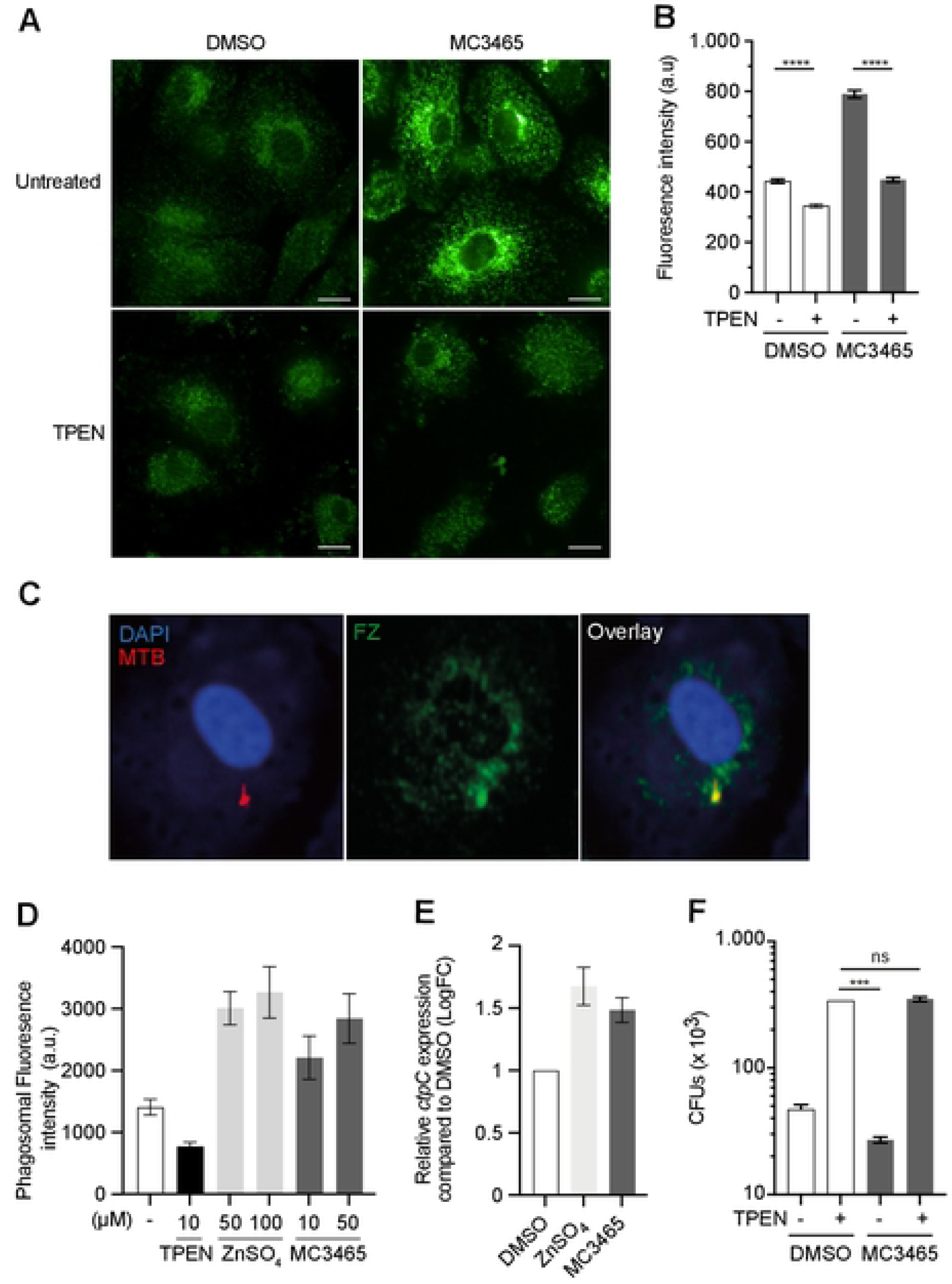
Free zinc is released upon MC3465 treatment and accumulates within the mycobacterial phagosome. **(A)** Macrophages were infected with MTB during 2 h and incubated with the zinc-chelating agent TPEN (2.5 µM). After 1 h, cells were treated with MC3465 for 4 h. Cells were fixed and stained with the free zinc-specific fluorescent probe FluoZin^TM^3-AM. Scale bar, 10 μm. **(B)** Average FluoZin^TM^3-AM signal intensity, for at least 500 cells per condition. **(C-D)** Macrophages were infected with DsRed-MTB and incubated with TPEN, ZNSO_4_ or MC3465 during 4 h. Cells were then fixed and stained with FluoZin^TM^3-AM. **(C)** Representative image of FluoZin^TM^3-AM colocalization with MTB. **(D)** Quantification of FluoZin^TM^3-AM colocalization with DsRed-MTB. At least 100 phagosomes were counted per condition. **(E)** RT-qPCR analysis of *ctpC* expression upon incubation with MC3465 or ZnSO_4_. After 4 h treatment, total cellular RNA was extracted and analyzed by RT-qPCR. Data are are normalized relative to *ftsZ* gene. One representative experiment (of at least two) is shown. **(F)** MTB-infected cells were incubated with TPEN. After 1 h, cells were treated with MC3465. The number of intracellular bacteria was enumerated at 96 h post-treatment. One-way ANOVA test was used. Error bars represent the mean ± SEM. ns: not significant, ***p<0.001, ****p<0.0001. Error bars represent the mean ± SD. t test was used. *p<0.05, **p<0.01, ****p<0.0001.

It has been shown that zinc exposure leads to rapid induction of the putative heavy metal efflux P1-type ATPase ctpC (Rv3270) (27). We thus analyzed by RT-qPCR the expression of *ctpC* in cells incubated with MC3465 or the positive control ZnSO_4_ during 4 h. The expression of *ctpC* was increased in response to both compounds (Fig 3E). Overall, these results indicated that MTB is exposed to a high concentration of zinc during treatment with MC3465.

We then tested whether MC3465 limited intracellular MTB growth through the influx of zinc into the phagosome. MTB-infected macrophages were treated with TPEN, which inhibited the increase in the Fluozin^TM^ 3-AM signal (Fig 3A and 3B). After 1 h of treatment, cells were incubated with MC3465 and the number of bacteria was determined by CFUs at 96 h post-treatment. As shown in Fig 3F, TPEN abolished the effect of MC3465, demonstrating that zinc release is required for better control of MTB upon MC3465 treatment.

### Structure-activity relationship study of MC3465 allowed the identification of MC3466, a more potent analog, effective against clinical strains of MTB

We then sought to develop a more effective, yet endowed with low toxicity compound such as MC3465. Structure-activity relationship (SAR) study is a key aspect of any drug design and development campaign (32). MC3465 is a 1,2,4-oxadiazole derivative bearing a 4-bromophenyl moiety at C3 and a 3-bromopropyl chain at C5. We tested 29 analogues of MC3465 (structures in S2 Table) for their ability to restrain MTB growth within human macrophages, as described in Figure 1. These compounds possess different substitutions on the phenyl moiety at C3 and alkyl, haloalkyl, alkoxy, alkylaryl, or carboxamide chains at the C5 position, with two of them also having central cores different from the original 1,2,4-oxadiazole one. Using high-content screening confocal microscopy, we identified active and inactive analogs of MC3465 (S2 Table). This approach allowed us to identify the presence of an electron-withdrawing group at the *para* position of the C3-phenyl ring as very important for compound activity. Furthermore, the 3-bromopropyl chain guaranteed the best compound activity, and switching to other kinds of side-chains was detrimental, with the exception of compound MC3617, which has a carboxamide on C5.

Interestingly, we identified MC3466 as the most effective molecule of this series, with higher potency than MC3465, and the capability to reduce the area of GFP-expressing bacteria by 69% compared to the control (Fig 4A). Although the structures of MC3466 and MC3465 are very similar, MC3466 differs from MC3465 for the presence of a methoxy group, rather than a bromine atom, at the *para* position of the C3-phenyl ring (Fig 4B). We confirmed this result by counting the number of bacteria inside macrophages treated with MC3466 for 96 h (Fig 4C). MC3465 and MC3466 decreased MTB growth by 50% and 67%, respectively. As MC3465, the analogue MC3466 did not affect cell viability and had no effect on the growth of MTB in liquid culture medium at 10 μM nor at a higher concentration (50 μM) (Fig 4D). We next assessed the half maximal inhibitory concentration (IC_50_) of both MC3465 and MC3466. Briefly, macrophages were infected with MTB, and treated with a range of concentration of MC3465 and MC3466 for 96 h. The IC_50_ was estimated using image-based analysis with dose-response curve (DRC) or CFU-based assay. We obtained similar results with these two methods, with estimated IC_50_ values for MC3465 of 4.5 or 5.9 μM and 2.5 or 3.2 μM for MC3466, respectively (Fig 4E and 4F). As described for MC3465, MC3466 induced an increase in free zinc in macrophages (Fig 4G and 4H). Inhibition of this release by TPEN cancels out the compound’s effects on intracellular MTB growth (Fig 4I).

**Fig 4.**
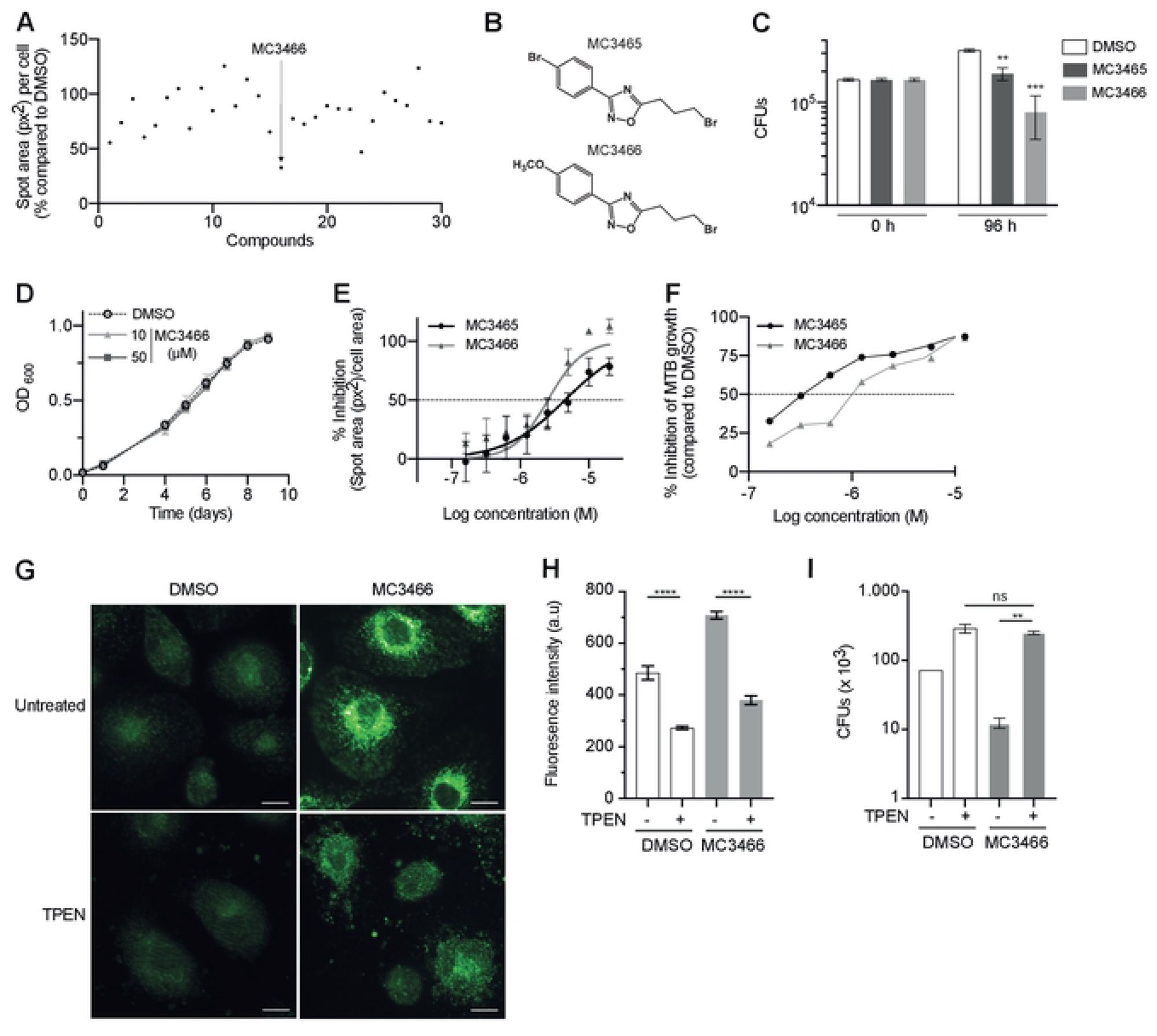
Structure-Activity Relationship (SAR) analysis of MC3465 and identification of MC3466 as a more effective analogue. **(A)** Macrophages were infected with GFP-MTB were incubated with 29 MC3465 analogues. 96 h post-infection, images were acquired by automated confocal microscopy followed by image analysis as described in Fig 1A and 1B. The spots area per cell was expressed as the percentage of GFP area in compound-treated cells compared to cells incubated with DMSO. **(B)** Structures of MC3465 and MC3466. **(C)** MTB-infected macrophages were treated with MC3465 or MC3466 (10 µM). After 96 h, the number of intracellular bacteria was enumerated. One representative experiment (of at least three) is shown. **(D)** MTB growth in liquid culture medium in the presence of the analogue MC3466 at different concentrations, determined by OD_600_. **(E)** The dose-response curves (DRC) for MC3465 and MC3466 were obtained by automated confocal microscopy followed by image analysis. The ratio of GFP area in compound-treated macrophages compared to cells incubated with DMSO, was normalized with the negative control DMSO (0% inhibition) and the positive control RIF (100% inhibition). **(F)** MTB-infected cells were treated with different concentration of MC3465 or MC3465. After 96 h, bacteria were enumerated by CFU and the IC50 of each compound was determined. **(G-I)** MTB-infected macrophages were incubated with TPEN. After 1 h, cells were treated with MC3466. **(G)** Representative image of FluoZin^TM^3-AM staining at 4 h post-treatment. **(H)** Average FluoZin^TM^3-AM signal intensity for at least 500 cells per condition. **(I)** The number of intracellular bacteria was enumerated at 96 h post-treatment. One-way ANOVA test was used. Error bars represent the mean ± SEM. ns: not significant, ***p<0.001, ****p<0.0001.

### MC3466 potentiates the activity of known anti-TB drugs

HDT aims to increase the success of TB treatment. However, HDT agents are more likely to be used in addition to conventional anti-TB drugs that directly target the bacteria (3). Our data have shown that MC3465 and MC3466 help human macrophages to control the growth of MTB but are insufficient to eradicate the bacteria. We therefore investigated whether MC3466 could act together with some anti-TB drugs to improve the efficacy of the treatment. We first tested the association between MC3466 and the anti-TB drugs, rifampicin (RIF) and bedaquiline (BDQ) in liquid culture medium. As expected, we showed no synergy between MC3466 and RIF or BDQ on the growth of MTB (S5A-C. Fig). We then focused on MTB-infected macrophages treated with MC3466 and different concentrations of the two drugs. After 96 h treatment, cells were lysed and bacteria counted. As expected, both anti-TB drugs alone induced a decrease in bacterial numbers compared to untreated cells in a dose dependent manner (Fig 5A and 5B). However, in the presence of MC3466, RIF and BDQ showed a higher bactericidal activity. At low concentration of the two antibiotics (0.1 µg/ml), MC3466 increased the efficiency of RIF by 93% and BDQ by 97%. This effect seems to go far beyond an additional effect of the compound and these anti-TB drugs; MC3466, RIF and BDQ alone reduced the bacterial load by 65, 65 and 79 % compared to DMSO respectively.

**Fig 5.**
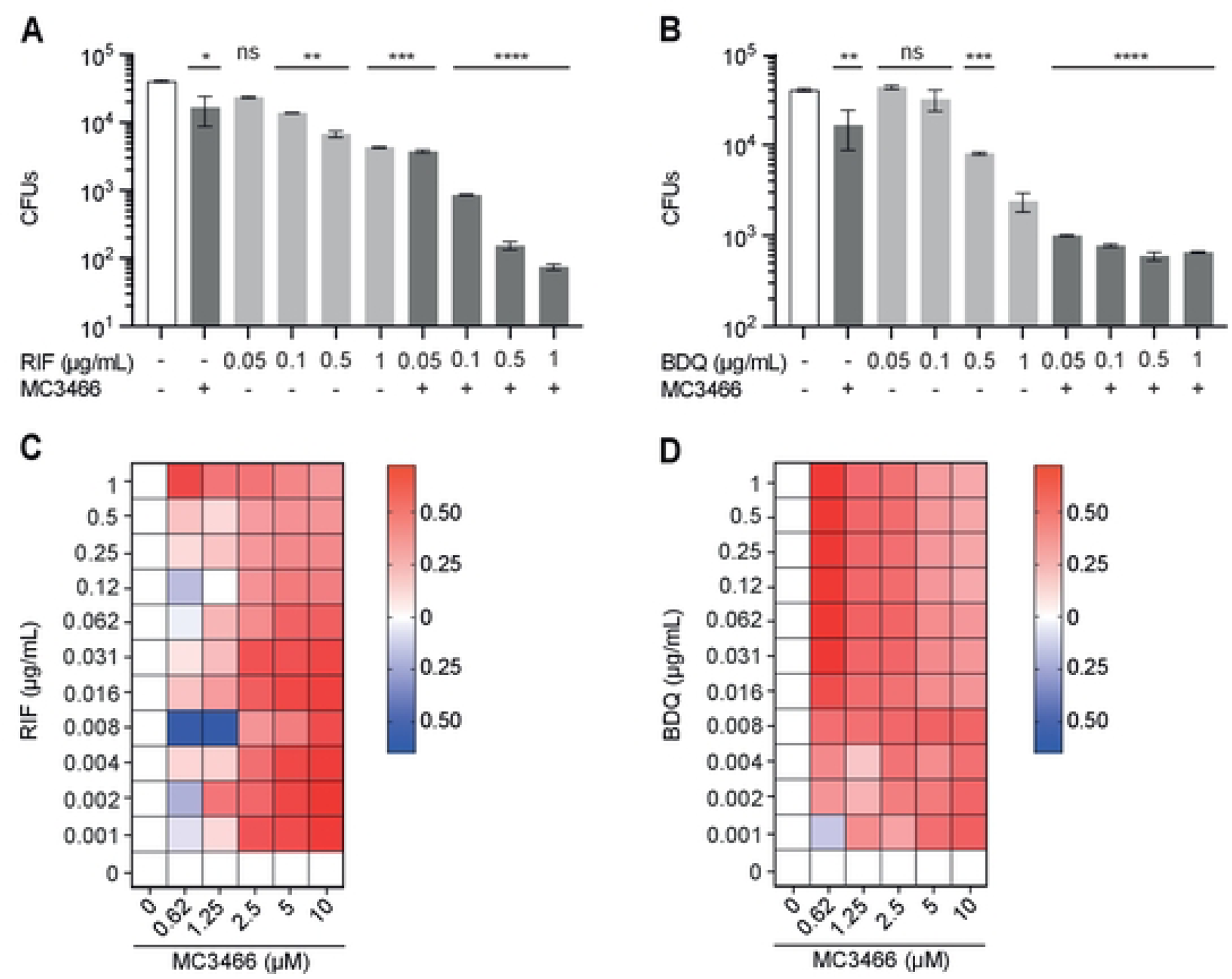
MC3466 potentialized the efficacy of rifampicin and bedaquiline in MTB-infected macrophages. **(A)** Macrophages were infected with the drug-susceptible MTB H37Rv and treated with various concentration of rifampicin (RIF) in association or not with MC3466. After 96 h, cells were lysed and the number of intracellular bacteria enumerated. **(B)** As in (A) except that RIF has been replaced by bedaquiline (BDQ). **(C-D)** Heatmaps representing variation in drug combination, using the Bliss independence model, ranging from antagonism (blue) to synergy (red). Cells were infected with GFP-MTB and were treated with a range of concentration of MC3466 and RIF or BDQ. 96 h post-treatment, images were acquired by automated confocal microscopy followed by image analysis as described in Fig 1. One representative experiment (of at least two) is shown. One-way ANOVA test was used. Error bars represent the mean ± SD.*p<0.05, **p<0.01, ***p<0.001, ****p<0.0001.

We next used the Bliss independence model to confirm these results. This model is widely used to determine independence, synergy, or antagonism between two compounds (33). The Bliss model of independence assumes independence between the combined molecules. It states that if two drugs have independent effects, then the effect of the combination should be equal to the product of the two single compounds. A positive Bliss score indicates synergy. Conversely, a negative Bliss score indicates antagonism. MTB-infected macrophages were treated with a wide concentration range of MC3466 alone or associated with different concentrations of RIF or BDQ. After 96 h treatment, intracellular GFP-expressing MTB growth was assessed by determining the GFP area per cell. We confirmed that the combination of MC3466 with RIF or BDQ was highly bactericidal on MTB. This effect was not the result of an additive effect between the two drugs, but of a synergy (Fig 5C and 5D).

### MC3466 potentiates the efficiency of RIF in the mouse model of TB

Many drugs with high *in vitro* efficacy fail to produce significant effects *in vivo*. Many factors affect their absorption, distribution or metabolism. Before testing our compounds *in vivo* in mice, we checked whether our molecules allowed murine macrophages to control MTB infection. Raw 264.7 cells (a murine macrophage cell line) were infected with MTB and treated with both MC3465 and MC3466. After 48 h, cells were lysed and bacteria were enumerated by CFU. Both molecules were equally active in murine and human cells (S6A Fig).

We next tested the toxicity of MC3465 and MC3466, in naive mice. We injected intraperitoneally MC3465 and MC3466, six days a week for two weeks in uninfected C57BL/6J mice. No loss of weight was observed during treatment (S6B Fig). After two weeks, total blood was collected to quantify the level of liver enzymes which might indicate inflammation or hepatocyte injury. No difference between the groups treated with the compounds and the control group was observed for the level of alanine aminotransferase (ALT) and aspartate aminotransferase (AST) (S6C and S6D Fig), suggesting there was no toxicity associated to both MC3465 and MC3466 treatments.

We then evaluated the efficacy of our compounds on MTB-infected mice in an acute model of infection, alone or in combination with RIF. C57BL/6J mice were infected via the aerosol route. After one week, mice were treated for two weeks, six days a week with MC3466 with or without RIF. Lungs were harvested, and the number of bacteria were estimated by CFUs (Fig 6A). The number of bacteria after the MC3466 treatment was decreased by 5 times. As observed *in vitro*, the combination of RIF and MC3466 was the most effective in decreasing the bacterial load (Fig 6A). The bacterial load was 31 times lower in MC3466/RIF-treated mice than in RIF-treated mice.

**Fig 6.**
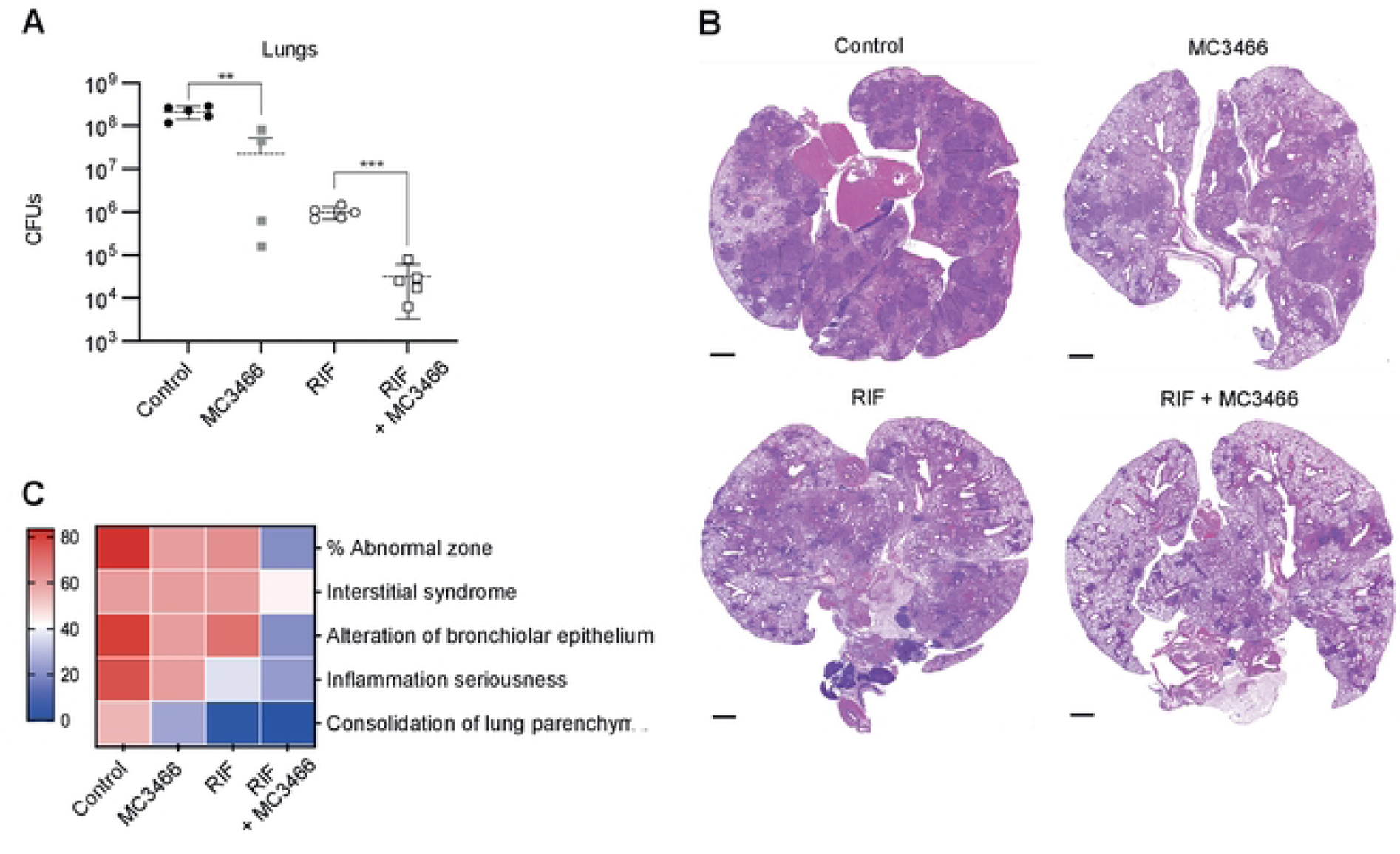
Synergy between RIF and MC3466 in MTB-infected mice. **(A)** C57BL/6J mice were infected by aerosol with MTB. After one week, mice were treated for two weeks, six days a week with MC3466 with or without RIF. Lungs were harvested, and the number of bacteria were estimated by CFU. **(B)** Representative hematoxylin and eosin stains of lungs, 2 weeks after treatment. Scale bar 1 mm. **(C)** Heatmap representing the histological scores upon RIF and MC3466 treatment. Lung sections are graded according to the percentage of abnormal area, presence of interstitial syndrome, alteration of bronchiolar epithelium, severity of inflammation, and consolidation of lung parenchyma. One representative experiment (of two) is shown. t test was used. Error bars represent the mean ± SD. t test was used. **p<0.01, ***p<0.001.

Histopathological observations of MTB-infected mice treated with RIF, MC3466 or both molecules confirmed these results. We observed large differences between the different groups, especially regarding the severity and organization of inflammatory lesions (Fig 6B and 6C). Lung sections of MC3466- or RIF-treated mice displayed significant interstitial syndrome and alteration of bronchiolar epithelium; lungs of RIF-treated mice were less inflammatory. However, lungs of mice treated with both MC3466 and RIF resembled the lungs of healthy mice, with only residual inflammation.

## Discussion

Widespread antimicrobial resistance poses a threat to public health that requires prompt and adaptable approaches. High-throughput screening has been extensively used to identify potential targets for antimycobacterial agents (34) or drug candidates that inhibit MTB growth (35, 36). However, this approach has rapidly showed some limitations as drugs inhibiting MTB growth in culture medium may be ineffective on intracellular MTB. For instance, host drug-metabolizing enzymes and transporters may influence antimicrobial pharmacokinetics and pharmacodynamics, thereby impacting their efficacy and/or toxicity (37). The development of resistance to these molecules is also only a matter of time. In contrast, HDT is instead less prone to bacterial resistance and could also be used to reduce disease severity and mortality. Increasing the natural resistance of macrophages, the primary cell target of MTB, is indeed a promising avenue in the context of TB. Here using high content imaging on human MTB-infected macrophages, we identified 1,2,4-oxadiazole based compounds, with no effect on MTB growth in liquid culture, but able to inhibit the intracellular mycobacterial growth inside the host in a concentration dependent manner. Interestingly, these compounds have no similarity to other published 1,2,4-oxadiazole derivatives with activity on MTB (38–40).

Although the exact host target of MC3465 remains elusive, our data suggest that MC3465’s mechanism of action is different from those previously described for HDTs. Indeed, most of the HDTs render macrophages less permissive to MTB (3, 4) by favoring phagosome-lysosome fusion, activating autophagy, or inducing antimicrobial peptides. Our data clearly showed that there is no activation of autophagy in presence of MC3465. However, we demonstrated that, upon treatment with MC3465, the expression of genes encoding metallothioneins was downregulated in macrophages and associated with a release of free zinc inside the cells and a relocation inside the phagosome, leading to MTB metal poisoning. This defense mechanism is commonly used by macrophages, overloading the phagosome with copper and zinc rendering the vacuolar environment unsuitable for bacterial survival (27, 28). In response, MTB has developed different strategies to protect itself from metal poisoning (41). For instance, following zinc intoxication from macrophages, it has been shown that MTB upregulates the expression of genes encoding heavy metal efflux P-type ATPases CtpC, CtpG, and CtpV (27). Our data shows that MTB indeed increases the expression of *ctpC* in MC3465-treated macrophages. Altogether, our results demonstrate a novel mechanism of action of MC3465 as an HDT, related to zinc poisoning of the bacterial pathogen.

MC3465-treated macrophages control the growth of the laboratory MTB strain H37Rv and more virulent clinical MTB isolates, but they are still permissive to *L. monocytogenes* or *S* Typhimurium infection. This result is likely due to the difference of lifestyle within the host cells between mycobacteria, *L. monocytogenes* and *S.* Typhimurium. Upon uptake by macrophages, *L. monocytogenes* is engulfed in a phagosome but rapidly escapes to the cytosol where it multiplies (42). *S.* Typhimurium resides in a late endosome-like vacuole characterized by projection of filaments (43). The maturation of the mycobacterial phagosome is arrested at an early stage of biogenesis by exclusion of the V-ATPase. Under certain circumstances, *S.* Typhimurium and MTB may also translocate into the cytosol (43). Because of their very different intracellular lifestyle, MTB, *L. monocytogenes* and *S.* Typhimurium are thus likely exposed differently to different zinc concentrations following MC3465 treatment. Moreover, MTB, *L. monocytogenes* and *S*. Typhimurium have different sensitivity to zinc as they have developed their own strategies to sequester essential transition metals and protect themselves from toxicity. As an example, *Salmonella* appears to be less sensitive to the increase in cellular zinc levels following infection, as zinc supplementation and inactivation of metallothioneins 1 and 2 (which increased free zinc levels in macrophages), favor *Salmonella* survival (44). Different intracellular lifestyles associated to different resistances to zinc poisoning would explain the specificity of MC3465 towards mycobacterial strains.

In the fight against TB, there is growing interest in the development of effective antibacterial drug combinations for better therapeutic results (45, 46). Experimental drug regimens have been tested *in vitro* and optimized for their use in *vivo* (47). Recently, drug combinations using an already approved drug with an adjuvant such as small molecules, have been exploited for the design of new drug regimens. These molecules are also called boosters, activators or enhancers depending on the study (48). For example, several 1,2,4-oxadiazoles were tested for their potency to boost the antibacterial activity of ethionamide (39, 49). HDT, associated to pathogen-targeted approaches, is a very promising approach for overcoming antimicrobial resistance. One of the striking findings of our study is the potentiation of classical anti-TB drugs, *in vitro* and *in vivo*, by the treatment by MC3465 or its analog MC3466. Indeed, we demonstrated that suboptimal concentrations of a first- and a second-line anti-TB drugs, RIF or BDQ, respectively, in combination with our compounds were more effective in preventing intracellular growth of MTB than those used alone. More importantly, this synergistic effect was also demonstrated *in vivo* for RIF in a mouse model of TB. In parallel, we also showed there is no weight loss nor indication of inflammation and hepatocyte injury, as the level of alanine aminotransferase (ALT) and aspartate aminotransferase (AST) remains stable following two weeks of treatment with our compounds.

One of the possible applications of HDT in TB is to limit inflammation and tissue damage (4). Histological analysis of lungs of MTB-infected mice treated with RIF, MC3466 or both, revealed that association of RIF with MC3466 considerably improves the healing of lesions. This is likely due to the greater efficacy of RIF when used in combination with MC3466 but may also be due to the downregulation of the type I IFN pathway upon treatment with our compound. Indeed, our transcriptomic analysis of MC3465-treated macrophages showed indeed that genes involved in the type I IFN pathway are also downregulated after 24h treatment. Several studies have demonstrated that type I IFN overexpression is deleterious for the host during MTB infection (50). *In vitro*, type I interferon and its downstream signaling cascade inhibited the antimicrobial response induced by type II interferons in human monocyte (51). In patients with active TB, blood transcriptional gene signature with overexpression of type I IFN-related genes have been correlated with disease severity and is downregulated following successful treatment (50, 52). Targeting the type I IFN pathway may be used as HDT. In the absence of IL-1, PGE2 failed to inhibit type I IFNs. The resulting over-expression of these cytokines leads to increased lung pathology. Administration of PGE2 and zileuton, a 5-LO inhibitor, negatively regulates type I IFN *in vivo* and confers protection in *Il1*α, *Il1*β^−/−^ mice infected with MTB (53).

In summary, we identified new compounds based on 1,2,4-oxadiazole scaffold which allow human macrophages to control MTB replication and to potentiate the efficacy of two potent anti-TB drugs, namely RIF and BDQ. These observations suggest that MC3465 and its analogs merit further investigation as potential HDTs.

## Materials and Methods

### Ethics Statement

Buffy coats were obtained from anonymous healthy donors who signed a consent to donate their blood for research purposes. The blood collection protocol was approved by the French Ministry of Research and by an ethics committee of the Etablissement Français du Sang (n°12/EFS/134). Animal studies were performed in agreement with European and French guidelines (Directive 86/609/CEE and Decree 87–848 of 19 October 1987). The study was approved by the Institut Pasteur Safety Committee (Protocol 11.245) and the ethical approval by local ethical committees “Comité d’Ethique en Experimentation Animale Institut Pasteur N° 89 (CETEA)” (CETEA dap 200037).

### Macrophages and cell lines

Blood mononuclear cells were isolated from buffy coats by Lymphocytes Separation Medium centrifugation (Eurobio). CD14+ monocytes were isolated by positive selection using CD14 microbeads (Miltenyi Biotec) and were cultured at 37°C and 5% CO_2_ in RPMI-1640 medium (Gibco) supplemented with 10% heat-inactivated fetal bovine serum (FBS; Dutscher), and 2 mM L-glutamine (Gibco) with macrophage colony stimulating factor (M-CSF, 20 ng/mL; Miltenyi Biotec) (hereafter defined as complete medium). After six days of differentiation, the resulting macrophages were incubated in a buffer solution containing PBS, 2 mM ethylenediaminetetraacetic acid (EDTA), at 37°C and 5% CO_2_ for 15 minutes before being harvested and counted.

THP-1 cell line was cultured in RPMI-1640 medium (Gibco) supplemented with 10% FBS and L-glutamine (Gibco). Mouse embryonic fibroblasts (MEF) and Henrietta Lacks (HeLa, ATCC, CCL-2) cells were cultured in Dulbecco’s Modified Essential Medium supplemented with L-glutamine (Gibco) and 10% FBS. Cells were incubated at 37°C, 5% CO_2_, and the culture medium was changed every two days.

### Bacterial strains

*Listeria monocytogenes* strain wildtype EGD, number BUG600 was grown in brain-heart infusion medium (BHI; Difco Laboratories) at 37°C. *Salmonella enterica* serovar Typhimurium (strain Keller) was grown in Luria Bertani (LB) medium. Mycobacteria were grown at 37°C in Middlebrook 7H9 medium (Becton-Dickinson) supplemented with 10% albumin dextrose catalase (ADC, Difco Laboratories), and 0.05% Tween 80. Hygromycin B was added to the culture medium of strains containing a plasmid encoding *gfp* or *dsred* gene. MTB strains used in this study include MTB strain H37Rv transformed with an Ms6-based integrative plasmid pNIP48 harboring GFP or DsRed protein (34), *M. bovis* BCG Pasteur, CDC1551, GC1237, Myc5750 and BDQ-resistant MTB strain H37Rv (54). Rifampicin resistant MTB strain was provided by Anne-Laure Roux and Jean-Louis Herrmann (Assistance Publique Hôpitaux de Paris, Hôpital Ambroise Paré, France).

### Construction of *Rv1151c* deletion mutant

ΔRv*1151*c knock-out strain was obtained by performing DNA recombination. Briefly, 750-bp fragments of the upstream and downstream regions of Rv*1151*c gene were amplified by PCR from H37Rv genomic DNA. Primers used for Rv*1151*c upstream region are 5’-tgatgtacctacaacccgaac-3’ and 5’-gactgagcctttcgttatttaaataatccgttcttgtcatcgcggaacgtcggt-3’. Primers used for Rv*1151*c downstream region are 5’-cgttccactgagcgatttaaattgatcgaagtcaatcccgagcccacgccg-3’ and 5’-ggtggattccacgaacgtgc-3’. The zeocin cassette was also amplified by PCR using primers 5’-atttaaataacgaaaggctcagtc-3’ and 5’-atttaaatcgctcagtggaacg-3’. A linear fragment composed of the zeocin cassette flanked by the Rv*1151*c upstream and downstream regions was amplified by PCR using 5’-aaaatatgatattcgcatggcg-3’ and 5’-aaagctcgaaagccgctggt-3’. This linear fragment was then electroporated into H37Rv harboring the pJV53-kanamycin plasmid, previously grown in 7H9 medium supplemented with 0.2% acetamide for 24h. After electroporation, 1 ml of 7H9 medium is added to the bacteria followed by an incubation step at 37°c for 48h. Transformants are then plated on 7H11 containing both kanamycin and zeocin antibiotics. After 3 weeks of incubation at 37°C, kanamycin-zeocin resistant clones were screened by PCR using 5’-actacagctggatggattccg-3’ and 5’-acacgccagcgtcagcaatc-3’.

### Bacterial infection

Before infection, MTB were washed and resuspended in 4 mL PBS. Clumps were eliminated by passing through a 10 µm filter. The density of bacteria in the supernatant was verified by measuring the OD_600_ and aliquot volumes defined to allow one bacterium-per two cell infections. After 2 h of incubation at 37 °C, infected cells were washed three times in PBS and treated with amikacin (50 µg/ml). After 1 h, cells were washed and incubated in fresh complete medium.

*L. monocytogenes* and *S.* Typhimurium were grown to exponential phase. Bacteria were washed twice in PBS and added to cells at a multiplicity of infection (MOI) of 50:1. After 1 h, cells were washed in PBS, and gentamycin (50 μg/ml) was added to kill extracellular bacteria. After 1 h, cells were washed and incubated in fresh complete medium.

### Compound synthesis

Compounds MC3465, MC3466, MC3209, MC3581, MC3582, MC3469, MC3453, MC3564, MC3565, MC3617, MC3459, MC3220, MC3212, MC4214, and MC4209 were synthesized as previously reported (15). Full details regarding the synthesis and physico-chemical characterization of compounds MC3618, MC3586, MC3610, MC3577, MC3579, MC3735, MC3738, MC3775, MC3750, MC3748, MC3903, MC3904, MC3905, MC3573, and MC3578, will be reported elsewhere. All new compounds had spectral (^1^H-NMR, ESI-MS) data in agreement with their chemical structures.

### Image-based high-content screening

Macrophages were infected with GFP-expressing H37Rv (MTB-GFP) and were cultured in 384-well plate (Cellcarrier plate, PerkinElmer); each well containing a compound resuspended in DMSO at final concentration of 10 μM. Each compound was tested in triplicates. DMSO and rifampicin (10 µg/mL) were used as negative and positive controls, respectively. 96 h post-infection, cells were fixed with 4% paraformaldehyde at room temperature (RT). After 1 h, cells were washed and stained with HCS CellMask Blue stain (2 μg/mL, Thermo Fisher) and Hoechst 33342 (5 μg/mL, Thermo Fisher) in PBS for 30 minutes at room temperature, washed twice with PBS and stored at 4 °C until acquisition. Images were acquired using the automated confocal microscope Opera Phenix High-Content Screening System (Perkin Elmer Technology) with a 40x/NA 1.1 water objective followed by image analysis (Columbus Conductor™ Database, Perkin Elmer Technologies). Data analysis was carried out to obtain numbers of cells, percentages of infected cells and quantified areas of intracellular bacteria (px^2^: pixels^2^).

### Cell viability assay

Cell viability was determined using the MTT assay kit (Abcam), according to manufacturer’s instructions.

### Resazurin assay determination of the minimal inhibitory concentration (MIC)

The microdilution test was performed in 96-well plates as previously described (54, 55). Briefly, MTB were cultured in 7H9 liquid medium containing 2-fold dilutions of antibiotics during 6 days. The dye resazurin (Sigma) at 0.02% (wt/vol) was then added to each well. After 24 h, the absorbance was measured at 570 nm.

### Determination of bacterial counts

Macrophages were lysed in distilled water with 0.1% Triton X-100. The number of bacteria was determined by plating ten-fold serial dilutions of the lysate, in triplicate, onto 7H11 agar plates. CFUs were scored after three weeks at 37°C. *L. monocytogenes* and *S.* Typhimurium were plated on BHI and Luria-Bertani agars, respectively. CFUs were counted after 24 h at 37°C.

### Indirect immunofluorescence

Macrophages were cultivated on coverslips in 24-well tissue culture plates for 24 h. Cells were infected with MTB-GFP and treated with MC3465. After 4 h and 48 h, cells were fixed with 4% paraformaldehyde for 1 hour at RT. LC3 labeling was performed as previously described (54). LC3B puncta were analyzed using a Leica TCS SP8 confocal microscope and quantified using ImageJ. Dot plots represent the mean values of at least 100 cells. Error bars depict the standard deviation.

### RNA isolation, library preparation and sequencing

Total RNA from macrophages was extracted using QIAzol lysis reagent (Qiagen) and purified over RNeasy columns (Qiagen). The quality of all samples was assessed with an Agilent 2100 bioanalyzer (Agilent Technologies) to verify RNA integrity. Only samples with good RNA yield and no RNA degradation (ratio of 28S to 18S, >1.7; RNA integrity number, >9) were used for further experiments. cDNA libraries were prepared with the Illumina TruSeq Stranded mRNA and the IDT for Illumina TruSeq UD Indexes and were sequenced on an Illumina NovaSeq 6000 systems at the CHU Sainte-Justine Integrated Centre for Pediatric Clinical Genomics (Montreal, Canada). Raw reads derived from the sequencing instrument are clipped for adapter sequence, trimmed for minimum quality (Q30) in 3’ and filtered for minimum length of 32 bp using Trimmomatic (56). Surviving read pairs were aligned to GRCh38 by the ultrafast universal RNA-seq aligner STAR (57) using the recommended two passes approach. Aligned RNA-Seq reads were assembled into transcripts and their relative abundance was estimated using Stringtie (58). All of the above processing steps were accomplished through the GenPipes framework (59). Exploratory analysis was conducted using various functions and packages from R and the Bioconductor project (60). Differential expression was conducted using both edgeR (61) and Limma (62).

### Data availability

The raw fastq files have been deposited in NCBI’s Gene Expression Omnibus (Edgar et al., 2002) and are accessible through GEO Series accession number GSE222412 (https://www.ncbi.nlm.nih.gov/geo/query/acc.cgi, token: ctazqyeavbcvnqp).

### Intracellular zinc labelling

Cells were fixed and stained with the cell permeant fluorescent FluoZin™-3, AM (Thermo Fisher; 2µg/mL) for 50 minutes in PBS at RT. Images were acquired with the automated fluorescence microscope Opera Phenix High-Content Screening System, with a 40x/NA 0.6 water objective. Fluorescence intensity was analyzed in Columbus Conductor on at least 400 different cells.

### Quantitative reverse transcription PCR (RT-qPCR)

Reverse transcription of mRNA to cDNA was done using SuperScript III Reverse Transcriptase (Thermo Fisher) followed by amplification of cDNA using Power SYBR Green PCR Master Mix (Thermo Fisher). Reactions were performed using a StepOnePlus Real-Time PCR System Thermal Cycling block (Applied Biosystems). The relative gene expression levels were assessed according to the 2^-ΔCt^ method (63).

The primers used in this study are: *ctpC* (*ctpC*-F: TCACCATTTTCACCGGGTAT; *ctpC*-R: GATGTTGAGCAACCACAGGA), *ftsZ* (ftsZ-F: CGGTATCGCTGATGGATGCTTT;ftsZ-R: CGGACATGATGCCCTTGACG), *gapdh* (*gapdh*-F: AATGAAGGGGTCATTGATGG; *gapdh*-R: AAGGTGAAGGTCGGAGTCAA) and *sirt2* (*sirt2*-F: TCACACTGCGTCAGCGCCAG; *sirt2*-R: GGGCTGCACCTGCAAGGAG).

### *In vivo* administration of MC3465 and MC3466

Seven-week-old female C57BL/6J mice were purchased from Charles River Laboratories and maintained according to Institut Pasteur guidelines for laboratory animal husbandry.

Compounds were solubilized daily in 0.5% DMSO + 4.5% Cremophor EL® + 90%NaCl. Mice were treated with MC3465 and MC3466 by intraperitoneal injections for 2 weeks, 6 days per weeks. Mice were deeply anesthetized with a cocktail of ketamine (Merial) and xylasine (Bayer) before cardiac puncture. Blood was collected in heparin-coated tubes. Aspartate aminotransferase (AST) and alanine aminotransferase (ALT) were quantified using the Reflover®Plus analyser.

Mice were infected via aerosol, generated from a suspension containing 5 × 10^6^ CFUs/ml of H37Rv to obtain an expected inhaled dose of 100–200 bacilli/lungs. One day after infection, two mice were killed, and the numbers of CFU were determined. After 7 days, mice were treated with MC3466 by intraperitoneal injections and/or with rifampicin (10 mg/kg) by gavage for 2 weeks, 6 days per weeks. CFUs in lungs of infected animals were determined after 21 days post infection.

### Histopathology

Lung samples were fixed in formalin for 7 days and embedded in paraffin. Paraffin sections (4-µm thick) were stained with hematoxylin and eosin (H&E). Slides were scanned using the AxioScan Z1 (Zeiss) system and images were analyzed with the Zen 2.6 software. Histopathological lesions were qualitatively described and when possible scored, using: (i) distribution qualifiers (i.e., focal, multifocal, locally extensive or diffuse), and (ii) a five-scale severity grade, i.e., 1: minimal, 2: mild, 3: moderate, 4: marked and 5: severe. For the histological heatmaps, the scores were determined as follows: the percentage of abnormal zone was estimated from low magnification images of scanned slides. All other scores were established at higher magnification (20–40× in the Zen program); the interstitial and alveolar syndrome scores reflected the extent of the syndrome, while the inflammation seriousness represented an evaluation of the intensity of the inflammatory reaction, i.e., abundance of inflammatory cells and exudate, conservation or disruption of the lung architecture; the bronchiolar epithelium alteration score was derived from both the extent and the severity of the lesions.

### Quantification and statistical analysis

Data were expressed as mean ± standard deviation (shown as error bar). Statistical analyses were performed with Prism software (GraphPad Software Inc), using the t test and one-way analysis of variance (ANOVA) as indicated in the figure legends. Differences between groups were examined statistically as indicated (*p < 0.05, **p < 0.01, ***p < 0.001 and ****p<0.0001). Results were considered statistically significant with a p-value < 0.05.

## Acknowledgments

We gratefully acknowledge the UTechS Cytometry and Biomarkers and the UTechS Photonic BioImaging (PBI) /C2RT of Institut Pasteur (Paris, France). PBI, part of the France–BioImaging infrastructure network (ANR-10–INSB–04; Investments for the Future) kindly acknowledge the support of the Région Île-de-France (DIM1 Health) for the use of the Opera Phenix. We also thank Charles Privé (CHU Sainte-Justine Integrated Centre for Pediatric Clinical Genomics, Montreal, Canada) for their technical support. AM was supported by the Fondation pour la Recherche Médicale (FRM FDM201806006250). D.R. acknowledges Sapienza Ateneo Project 2021 RM12117A61C811CE and Regione Lazio PROGETTI DI GRUPPI DI RICERCA 2020, project ID: A0375-2020-36597. An. M acknowledges the project FISR2019_00374 MeDyCa. This research project was financed by Institut Pasteur, the Georges, Jacques and Elias Canetti Award and the French National Research Agency (ANR Mustart 20-PAMR-0005). The text represents the authors’ views. The funders had no role in the study design, data collection or analysis, the decision to publish, or the preparation of the manuscript.

## Author Contributions

A.M. and L.T. conceived and designed all experiments. A.M. performed most of the experiments with the help of L.T.. C.K. validated the *in vitro* efficacy of some compounds. F.F., An.M. and D.R. provided the compound library and prepared the MC3465 analogs. A.D., N.A. and P.B. participated in the compound library screening. E.L. generated the ΔRV1151c mutant. M.E. under the supervision of M.H. provided SIRT2 deficient cells. A.P. and W.F. participated in the mouse experiments. D.H. performed the histological examination. S.G., A.G.G. and L.B. did the transcriptomic analysis. R.B. supervised the project; A.M. and L.T. wrote the original manuscript draft with the help of E.L.. All authors approved the final manuscript.

## Supporting information

**S1 Table. Epigenetics compound library used in this study.**

**S2 Table. List and structure of MC3465 analogs.**

**S1 Fig. MC3465 is non-toxic and limits intracellular growth of MTB, independently of macrophage polarization.**

**(A)** Naïve macrophages were treated with different concentration of MC3465. After 96 h, cell viability was assessed using the MTT assay according to the manufacturer’s instructions. **(B)** Human monocytes were differentiated with granulocyte Mϕ colony-stimulating factor (GM-CSF) or Mϕ colony-stimulating factor (M-CSF) for 6 days. Cells were then infected with MTB and treated with MC3465 for an additional 4 days. Cells were lysed and bacteria were enumerated by CFU. One representative experiment (of at least two) is shown. Error bars represent the mean ± SD. t-test was used. *p<0.05, **p<0.01.

**S2 Fig. MC3465 constrained the intracellular growth of MTB independently of SIRT2.**

**(A)** MTB-infected macrophages were treated with different concentrations of MC3465 and of two SIRT2 inhibitors, namely AGK2 and SirReal2. After 96 h, cells were lysed and bacteria were enumerated by CFU. **(B-C)** *SIRT2* expression was downregulated in macrophages using siRNA-mediated gene silencing. **(B)** Relative *SIRT2* expression measured by RT-qPCR in SIRT2-silenced cells. Relative expression levels were normalized to the *GADPH* gene. Scramble siRNA (scRNA) was used as a negative control. **(C)** Wild-type (WT) and SIRT2-silenced macrophages were infected with MTB and treated with MC3465 (10 µM). After 96 h, the number of intracellular bacteria was counted. **(D)** WT mouse embryonic fibroblasts (MEFs) or SIRT2^−/−^ MEFs were infected with MTB and treated with MC3465. After 48 h, cells were lysed and the number of intracellular bacteria was enumerated. **(E)** Growth of MTB H37Rv or RvΔ1151c in liquid culture medium, determined by OD600. **(F)** Human monocyte-derived macrophages were infected with either MTB wild-type (H37Rv) or a Rv1151c null mutant (ΔRv*1151c*). Cells were then treated with MC3465 and MC3466. After 0 or 96 h, bacteria were enumerated by CFU. One representative experiment (of at least two) is shown. One-way ANOVA test and t-test were used. Error bars represent the mean ± SD. *p<0.05, **p<0.01, ***p<0.001.

**S3 Fig. MC3465 did not induce autophagy in MTB-infected macrophages.**

**(A)** Detection by indirect immunofluorescence of LC3 (red) in MTB (green) infected macrophages, treated with MC3465 for 4 h (scale bar: 10 μm). DAPI (blue) was used to visualize nuclei. **(B)** Determination of the number of LC3-positive puncta per cell (unpaired t-test) after 4 h and 48 h treatment with MC3465. **(C)** MTB-infected macrophages were left untreated or incubated with MC3465 plus bafilomycin (BAF), 3-methyladenine (3-MA) or chloroquine (CQ). After four days, the number of intracellular bacteria was enumerated. One representative experiment (of at least two) is shown. Error bars represent the mean ± SD. One-way ANOVA test was used. ns: not significant, *** p < 0.001.

**S4 Fig. Free zinc is released upon MC3465 treatment.**

Naïve macrophages were incubated with ZNSO_4_, MC3465 or the zinc-chelating agent TPEN during 4 h. **(A)** Cells were fixed and stained with FluoZin^TM^3-AM. Scale bar, 20 μm. **(B)** Average FluoZin^TM^3-AM signal intensity, for at least 500 cells per condition.

**S5 Fig. Susceptibility and resistance of MTB strains to MC3465, MC3466, RIF and BDQ.**

**(A-C)** WT MTB H37Rv was incubated in liquid culture medium with different concentrations of MC3465 or MC3466 **(A)**, RIF **(B)** or BDQ **(C)** alone or in association with MC3465 or MC3466. The growth was determined by OD_600_. One representative experiment (of at least three) is shown. Error bars represent the mean ± SD.

**S6 Fig. MC3465 and MC3466 were well tolerated in mice.**

**(A)** The murine macrophage cell line, RAW 264.7, was infected with MTB and treated with MC3465 or MC3466. After 0 or 48 h, bacteria were enumerated by CFU. **(B-D)** Eight-week-old female C57BL/6J mice received different concentration of MC3465 and MC3466 by intraperitoneal injection 6 days per week during two weeks. **(B)** Weight was evaluated before each injection. **(C-D)** Blood was drawn before mouse sacrifice. **(C)** Alanine aminotransferase (ALT) and **(D)** aspartate aminotransferase (AST) levels were measured according to the manufacturer instructions. Error bars represent the mean ± SD. t test was used. ns: not significant, * p < 0.05, ** p < 0.01.

